# Behavioral Thermoceptive Responses and Morphologic Correlates in Mouse Models of CMT1A, HNPP and Aging

**DOI:** 10.1101/2025.11.11.686785

**Authors:** Vaibhav Oberoi, James O. Campbell, Nelson Akpabli-Tsigbe, Fatima Imran, Silvano Bond, Sara Ngo Tenlep, Charles D. Brennan, Meifang Wang, Smita Saxena, Kathryn R. Moss, De-Pei Li, W. David Arnold, Ryan Castoro

## Abstract

**Background:** Thermoceptive dysfunction is a frequent but understudied feature of peripheral neuropathies and aging. Patients often report abnormal heat perception, yet the underlying sensory mechanisms remain unclear. This study evaluated thermoceptive behavior and corresponding structural changes in mouse models of inherited dysmyelinating neuropathy and natural aging to identify shared and divergent mechanisms.

**Methods:** Thermal preference was assessed using a user-independent gradient apparatus spanning physiological to noxious temperatures, with automated quantification of time in zone, distance traveled, and velocity. Nocifensive responses were evaluated by hot plate latency. Intraepidermal nerve fiber density (IENFD) was measured in paw pads, and TRPV1-positive dorsal root ganglion (DRG) neurons were analyzed by immunofluorescence and confocal imaging.

**Results:** Thermal gradient testing revealed preserved temperature preference in CMT1A and HNPP mice but significantly altered behavior in aged animals, which spent less time in warmer zones. Hot plate testing showed prolonged times to nocifensive behavior in aged and CMT1A mice, whereas HNPP mice exhibited variable responses. IENFD was markedly reduced in aged mice but preserved in CMT1A and HNPP. DRG analysis revealed smaller soma diameters and reduced proportions of TRPV1-positive Aδ neurons in aged mice, while CMT1A animals maintained normal morphology.

**Interpretation:** Aging produces thermoceptive deficits through axonal degeneration and selective Aδ-fiber vulnerability, whereas CMT1A mice display conduction-related impairment due to dysmyelination. Both models reproduce key human sensory phenotypes and provide translational platforms for studying small-fiber dysfunction and therapeutic interventions in peripheral neuropathies.

**Key Points:**

- Thermoceptive dysfunction is a common but understudied feature of peripheral neuropathy and aging.
- In aged mice, altered thermal preference, reduced intraepidermal nerve fiber density, and smaller TRPV1-positive Aδ neurons indicate a degenerative mechanism of sensory loss.
- In CMT1A mice, delayed noxious heat responses occur despite preserved epidermal and DRG morphology, consistent with dysmyelination rather than axonal degeneration.
- HNPP mice showed inconsistent nocifensive responses without structural fiber loss, suggesting strain- or injury-dependent effects.
- Thermal gradient and hot plate assays demonstrate strong reproducibility and provide relevant outcome measures for preclinical modeling of thermoceptive dysfunction.

## INTRODUCTION

Temperature sensation, or thermoception, is a highly conserved neural function across both vertebrates and invertebrates(*1*). Peripheral thermoceptive signals are transduced by free, unencapsulated nerve endings in the skin and relayed to the dorsal horn of the spinal cord, where they synapse on second-order neurons(*2*). This occurs through two physiologically distinct classes of sensory fibers: unmyelinated C fibers, which convey slow, persistent changes in temperature(*3*), and thinly myelinated Aδ fibers, which provide rapid signaling, particularly in response to noxious heat(*4*). In peripheral neuropathies and age-associated neurodegeneration, thermoceptive function is frequently impaired and often associated with increased pain sensitivity, although the mechanisms remain poorly understood(*5, 6*). Critically, no therapies directly target the reduction in thermal perception, a common symptom in neuropathic diseases and preclinical characterization of thermoceptive deficits in inherited and aging models remains limited.

Charcot–Marie–Tooth disease (CMT) encompasses a heterogeneous group of monoallelic inherited neuropathies affecting both motor and sensory systems. Among more than 100 described variants, the most common is CMT1A, caused by a duplication of chromosome 17p12 containing the *PMP22* gene, resulting in abnormal myelin formation (dysmyelination)(*7, 8*). Clinical studies have reported that approximately 58% of CMT1A patients exhibit impaired Aδ fiber–mediated thermoception, a deficit that strongly correlates with neuropathic pain severity(*6*). This finding is particularly notable because epidermal nerve fiber densities are preserved in CMT1A(*9*), suggesting a predominantly functional rather than structural abnormality of Aδ fibers. Whether the phenotype of normal intraepidermal fiber counts with reduced noxious thermal responses also occurs in the widely used C3-PMP22 mouse model of CMT1A, which harbors additional copies of the human *PMP22* gene, is not known(*10*).

Hereditary neuropathy with liability to pressure palsies (HNPP) is another common inherited neuropathy resulting from a heterozygous deletion of chromosome 17p12 and consequent reduction in PMP22 expression(*7*). While patients are classically symptomatic after mechanical stress or compression, a subset develops persistent sensory polyneuropathy with impaired thermoception(*11*). However, the extent of Aδ fiber dysfunction and intraepidermal nerve fiber loss in HNPP remains poorly defined. The LacZ-PMP22KO mouse model of HNPP exhibits a severe dysmyelinating neuropathy at early post-natal ages similar to that what has been reported C3-PMP22 mice(*12*).

Age-related peripheral nerve degeneration represents a third major context for thermoceptive dysfunction. This process is heterogeneous, reflecting both intrinsic peripheral nerve aging (*13*) and systemic influences(*14*). In mice, motor axon degeneration and neuromuscular junction instability typically emerge by 20 months of age(*15*), whereas sensory deficits may appear as early as 11-12 months in some modalities and are associated with impaired dorsal root ganglion (DRG) electrophysiology and abnormal behavioral responses(*16, 17*). Aged rodents display blunted nociceptive responses to thermal stimuli, particularly noxious heat, but their behavioral responses to innocuous temperatures remain incompletely characterized(*17*). In humans, aging is associated with small-fiber loss in skin, although this is not consistently mirrored in sural nerve biopsies, highlighting the complexity of peripheral nerve aging.

Taken together, these observations underscore the need for systematic preclinical evaluation of thermoception across inherited and age-related neuropathies. In this study, we sought to characterize thermoceptive behaviors and correlate them with structural changes in small epidermal fibers and DRG sensory neurons in two inherited dysmyelinating neuropathy models (C3-PMP22 and LacZ-PMP22KO) and in naturally aged mice.

## METHODS

### 2.1 Animals

All experiments were conducted under protocols approved by the University of Missouri Institutional Animal Care and Use Committee (IACUC Protocol #46390). Mice were maintained under controlled conditions with a 12-hour light/dark cycle and ad libitum access to food and water. The LacZ-HNPP line, provided by Regeneron via Dr. Lucia Notterpek(*18*), was maintained on a 129S background and bred in the heterozygous state. The C3-PMP22 line, which carries additional copies of the human *PMP22* gene, was maintained on a C57BL/6J background (Jackson Laboratories) and also bred in heterozygosity. For aging studies, wild-type C57BL/6J mice were obtained at 20 months of age from Jackson Laboratories.

### 2.2 Thermal Gradient Testing

Behavioral testing was performed between 08:00 and 12:00 as done in other studies of thermoception(*19*). To minimize stress and ensure acclimatization, mice were habituated to the procedure room for one hour on the day prior to testing, followed by one hour of free exploration on the thermal gradient apparatus with the gradient turned off. The ambient temperature of the testing room was adjusted to 25°C and maintained throughout both habituation and testing.

On the test day, animals were again habituated at 25°C for 1 hour before being placed on the gradient device with the hot plate at 0 °C and 60 °C creating temperature zones (Figure 1A). Animals were placed onto the apparatus at the 23°C zone (Figure 1A), and behavior was recorded using an overhead GigE camera provided with the device (Imagesource, Germany) placed at 40 inches directly above the center of the device. Proprietary software provided with the device (Bio-Gradient version 2.0.46, BIOSEB, France) was used to adjust camera contrast settings within each lane and quantified the time spent in each temperature zone, the distance traveled in each zone, and the total distance traveled. It has been well described that thermoregulatory stabilization on gradient devices requires an initial adaptation period, only the final 15 minutes of the 60-minute run were analyzed, given prior work demonstrating body temperature habituation on similar devices occurs within the first 30 minutes(*20*).

**Figure 1.**
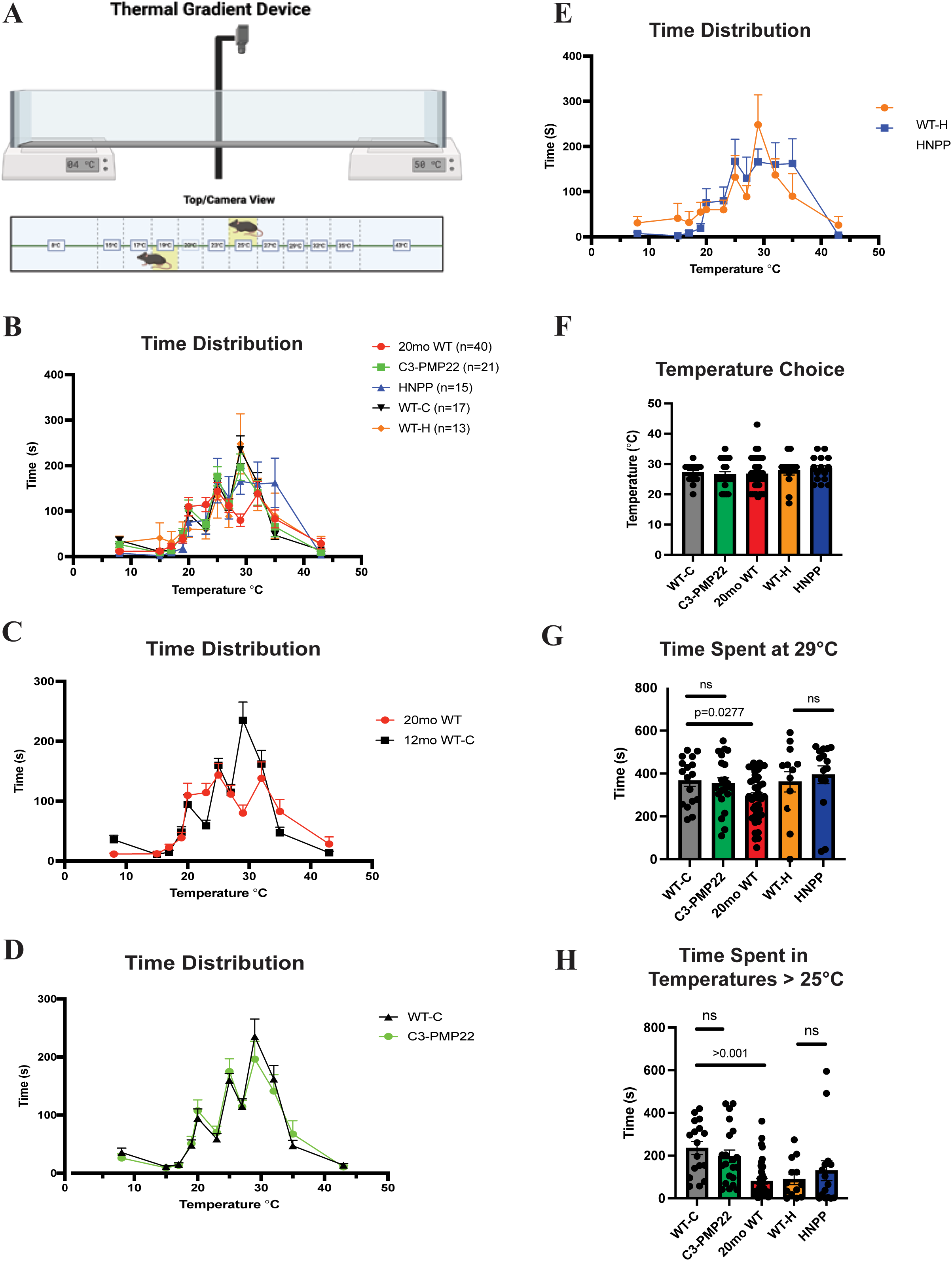
Thermal gradient assay and cumulative temperature preference behaviors. (A) Illustration of the thermal gradient device used in this study, created with BioRender (Castoro, R. [2025], https://BioRender.com/0p4wy0p). (B) Cumulative temperature preference behavior of all animals tested and by phenotype: 20-month-old wild type (C), CMT1A (D), and HNPP (E). (F) Comparison of cumulative temperature behavior between 20-month-old and 6-month-old wild-type mice. (G) Time spent at 29 °C, and (H) time spent at <25 °C for all animals. Data are presented as mean ± SEM. WT-C = 9-month-old wild-type littermates of C3-PMP22 mice (CMT1A, n = 21); WT-H = 4-month-old wild-type (129S) littermates of LacZ-HNPP mice (HNPP, n = 15); 20mo WT = 20-month-old wild-type (C57BL/6J, n = 40). One-way ANOVA followed by Bonferroni post hoc test was used for panels B–E; Student’s t-test was used for comparisons in panels F–H.

### 2.3 Hot Plate Assay

Thermal nociception was evaluated using a hot plate paradigm. On the day prior to testing, mice were habituated to the room for 1 hour and then placed on a room-temperature plate (25 °C) for 30 seconds, repeated three times with 5-minute intervals between trials. On the following day, animals were placed in a clear chamber on a plate preheated to 53 °C. The latency to the first nocifensive response, paw licking, hind limb elevation, or jumping, was recorded, and animals were immediately removed from the plate. Each mouse underwent four exposures: one acclimation trial followed by three test trials separated by 15-minute intervals. To prevent thermal injury, animals were removed if no response occurred within 30 seconds.

### 2.4 Reliability Analyses

The thermal gradient device version 2 (BIOSEB, France) is marketed as a user-independent system, and prior studies have confirmed low inter-rater variability. For this reason, inter-rater reliability was not formally reassessed. To evaluate intra-rater reliability, tested five wildtype C57BL/6J mice and five CMT1A mice in repeated trials spaced 4 weeks apart. Zone-specific time and distance traveled were compared across sessions to assess reproducibility.

Because hot plate testing is user-dependent and subject to greater variability, we assessed both intra- and inter-rater reliability. In the first cohort, consisting of five CMT1A and five wild-type mice, reproducibility was evaluated by repeated testing with two independent raters separated by one month(inter-rater) and a third separate evaluator (intra-rater). In a second cohort of twelve CMT1A and six wild-type mice, we assessed whether genotype-dependent differences could be reliably reproduced under standard testing conditions.

### 2.5 Intraepidermal Nerve Fiber Density

The day following testing euthanasia was performed with cervical dislocation while under isoflurane, paw pads were dissected from the right hind limb. Tissues were fixed for 2 hours at room temperature in 2% paraformaldehyde and cryoprotected overnight in 30% sucrose. Samples were embedded in OCT medium, frozen at −80 °C and sectioned at 40 µm using a cryostat. Sections were immediately washed in warm PBS to dissolve the OCT, followed by two additional PBS washes.

Permeabilization was performed with 0.1% Triton X-100 in PBS for 30 minutes, after which sections were blocked for 30 minutes in 3% bovine serum albumin with 0.05% Tween-20. Primary antibody incubation with anti-PGP9.5 (1:500; Cell Signaling Technology) was carried out for 48 hours at 4 °C in blocking buffer with gentle rocking. Sections were then washed and incubated with goat anti-rabbit Alexa Fluor 488 (1:2000; Jackson ImmunoResearch) for 1 hour at room temperature, counterstained with DAPI, and mounted on slides.

Images were acquired using an Olympus FV4000 confocal microscope with 20× and 60× objectives. Z-stacks spanning the full 40 µm thickness were collected and are represented by maximum intensity projections shown in figure 3A-E. Only fibers that penetrated the dermal–epidermal junction and extended into the nucleated epidermis were counted, following established criteria(*21*). For each animal, three nonadjacent sections covering approximately 3.0 mm of tissue were analyzed.

### 2.6 Dorsal Root Ganglion (DRG) Immunohistochemistry

Immediately after euthanasia, spinal columns were removed, and laminectomy was performed *in situ* to expose lumbar dorsal root ganglia (DRGs). DRGs were dissected, fixed overnight in 10% formalin, dehydrated through graded ethanol washes, and embedded in paraffin. Serial 10 µm sections were cut, with three consecutive slices mounted per slide. Antigen retrieval was performed for 30 minutes in citrate buffer (pH 6.0) using a pressure cooker at high temperature and pressure.

Sections were blocked overnight at 4 °C with a mixture of donkey, alpaca, and bovine sera (Jackson ImmunoResearch) in PBS containing 0.025% Tween-20. Primary antibodies included neurofilament heavy chain (NEFH, Invitrogen; 1:1250), phospho-ERK1/2 (Thr185/Tyr187; ThermoFisher; 1:100), TRPV1(DSHB; 1:50), and TRKA (FabGennix; 1:100). Antibody incubation was performed overnight at 4 °C.

After washing, sections were incubated with secondary antibodies (Jackson ImmunoResearch) consisting of donkey anti-mouse FITC (1:250), donkey anti-chicken Alexa Fluor 405 (1:500), bovine anti-goat Alexa Fluor 647 (1:3000), and alpaca anti-rabbit TRITC (1:500). Confocal imaging was performed using the same system and acquisition parameters as for paw pad tissue. Representative images were generated as maximum-intensity projections of z-stacks using Fluoview software.

The total number of neuronal cell bodies were counted under differential interference contrast microscopy (DIC) at both 20 and 60X. Quantification was performed only on sections containing 100 or more neuronal cell bodies. The percentage of all TRPV1⁺ was calculated as: (n = total number of TRPV1 stained cells / n = total number of neurons). Percentage Aδ neurons were calculated as: (n=TRPV1⁺/NEFH⁺/TRKA⁺) / (n = total number of TRPV1 stained cells) The cross sectional area of the neuronal cell bodies was measured using ImageJ.

### 2.7 Statistical Analysis

All statistical analyses were performed using GraphPad Prism (v10.0). Data were tested for normality using the Shapiro–Wilk test prior to parametric comparisons. Results are expressed as mean ± standard error of the mean (SEM). Comparisons among three or more groups were analyzed using one-way analysis of variance (ANOVA) followed by Bonferroni post hoc correction for multiple comparisons. Unpaired two-tailed Student’s t-tests were used for direct comparisons between two groups.

Pearson’s correlation coefficients were calculated to assess the relationship between hot plate latency times and soma cross-sectional area of TRPV1⁺ neurons. Intra- and inter-rater reliability for behavioral assays were evaluated using correlation analysis of repeated measures performed by blinded investigators.

## RESULTS

### 3.1 Thermal Gradient: Temperature Preferences and Zone Occupancy

Because sex-specific differences in thermal and nociceptive responses have been reported in some models, we first compared male and female mice within each genotype. As no significant sex differences were observed, and data were therefore combined for subsequent analyses (not shown).

When comparing thermal zone occupancy across experimental groups, older animals spent significantly less time in zones above 25°C compared with young controls, indicating an altered thermal preference (Figure 1B,F,G). In contrast, CMT1A (Figure 1C) and HNPP (Figure 1D) mice did not show differences from their respective wild-type littermates. Analysis of the preferred thermal zone, defined as the zone in which animals spent the longest duration, revealed no significant differences among genotypes or age-related difference (Figure 1E).

### 3.2 Thermal Gradient: Distance Traveled and Velocity

To evaluate whether locomotor capacity influenced thermal gradient behavior, we next analyzed distance traveled within each thermal zone and total distance across groups. Mean zone-specific distances were comparable among genotypes and aged mice groups (Figure 2A). However, when comparing total distance, aged wild-type and HNPP mice traveled significantly shorter distances relative to young wild-type controls, consistent with either reduced endurance or exploratory drive. CMT1A mice traveled distances were similar to wild-type controls (Figure 2B).

**Figure 2.**
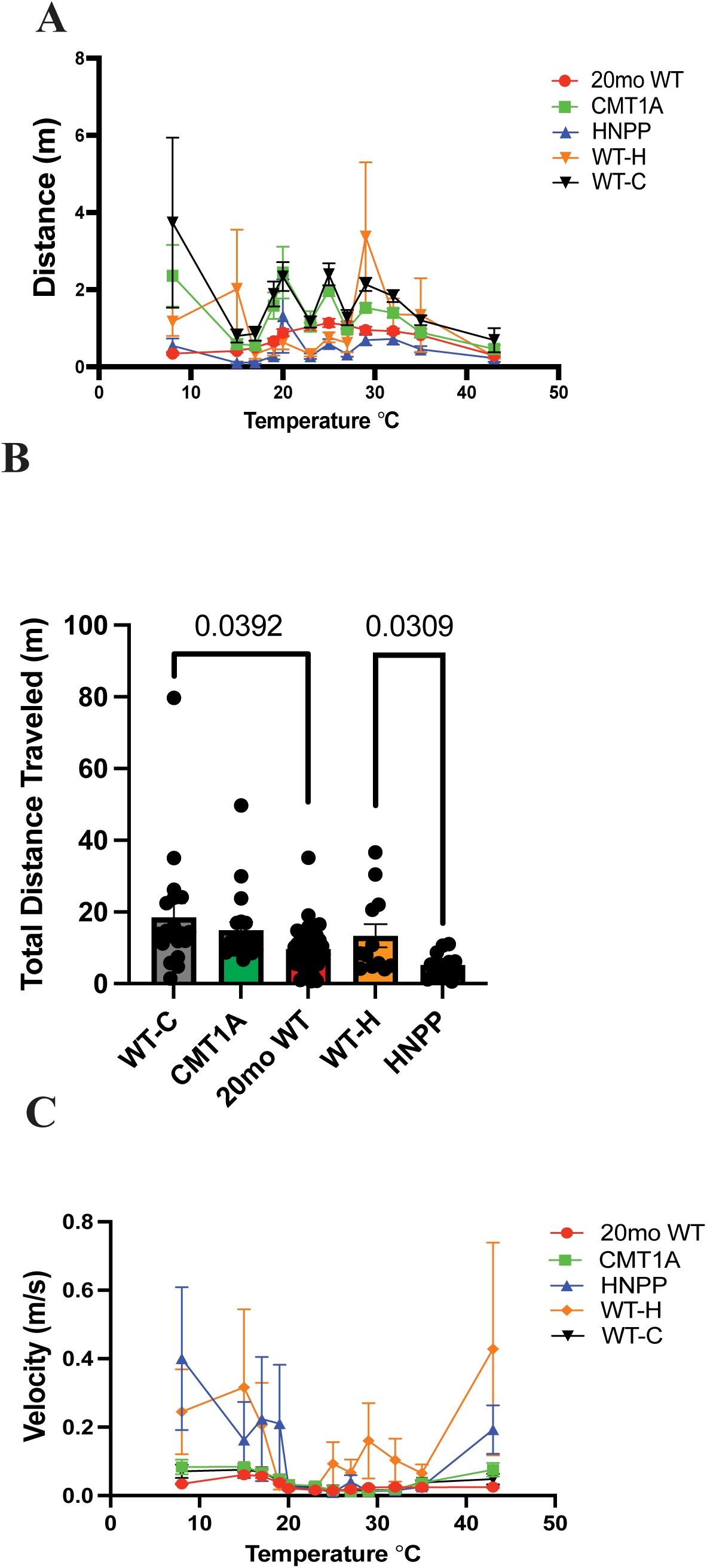
Locomotor performance during the thermal gradient assay. (A) Mean total distance traveled among all animals. (B) Total distance traveled during the final 15 minutes of the assay. (C) Average velocity per temperature zone for all animals tested. WT-C (n = 21) = 9-month-old wild-type littermates of C3-PMP22 (CMT1A, n = 21); WT-H = 4-month-old wild-type (129S) littermates of LacZ-HNPP (HNPP, n = 15); 20mo WT = 20-month-old wild-type (C57BL/6J, n = 40). One-way ANOVA followed by Bonferroni post hoc test was used for panels A and C; Student’s t-test was used for group comparisons in panel B.

Velocity was further examined to assess behavioral responses to noxious cold and heat. Across all groups, mice exhibited increased velocity when traversing noxious zones, consistent with an aversive response. However, no genotype- or age-dependent differences in velocity were detected (Figure 2C).

### 3.3 Hot Plate Responses

Nocifensive behaviors in the hot plate assay included paw licking, hindlimb lifting, and jumping, all of which could potentially be confounded by motor weakness. In preliminary tests across all groups at 4°C and 53°C, wild-type controls, CMT1A, and aged mice reliably exhibited multiple nocifensive behaviors, including jumping, indicating preserved motor capacity (Figure 3A). In contrast, wild-type 129S (WT-H) and HNPP mice did not exhibit consistent nocifensive responses even at 53°C, often reaching the 30-second cutoff without withdrawal, thereby increasing their risk of thermal injury. Expanded cohorts of 9-month-old CMT1A and aged mice confirmed significantly prolonged withdrawal latencies relative to wild-type controls (Figure 2B), consistent with impaired Aδ fiber–mediated nociception.

**Figure 3.**
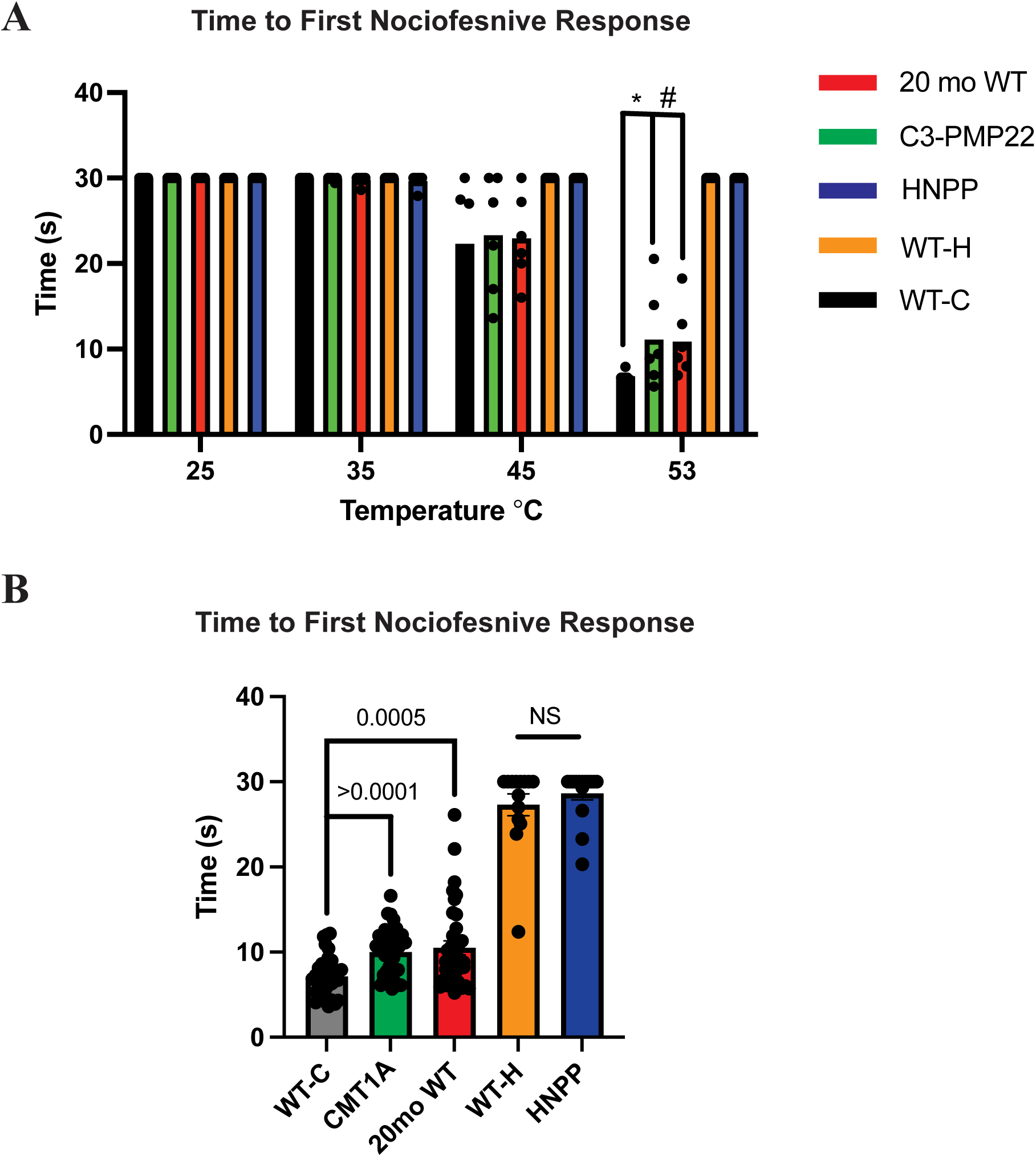
Hot plate testing across phenotypes. (A) Latency to nocifensive response across increasing temperatures in each phenotype tested n = 6 mice per phenotype. (B) Response latency at 53 °C in CMT1A (n =40), HNPP (n = 15), and 20-month-old wild-type (n=40) mice, WT-C (n = 40) WT-H (n = 13). Data are presented as mean ± SEM. *p = 0.1260; #p = 0.0621. Statistical comparisons were made using Student’s t-test; p > 0.05 was considered not significant.

### 3.4 Reliability of Thermal Gradient and Hot Plate Assays

To assess reproducibility, thermal gradient testing was repeated in the same cohort after one month. Cumulative temperature preference profiles remained highly consistent across sessions (Figure 4A–B). Zone-specific distances also showed similar distributions, with only minor reductions in distance traveled in cooler zones during the second run, while the total distance remained unchanged (Figure 4C–D).

**Figure 4.**
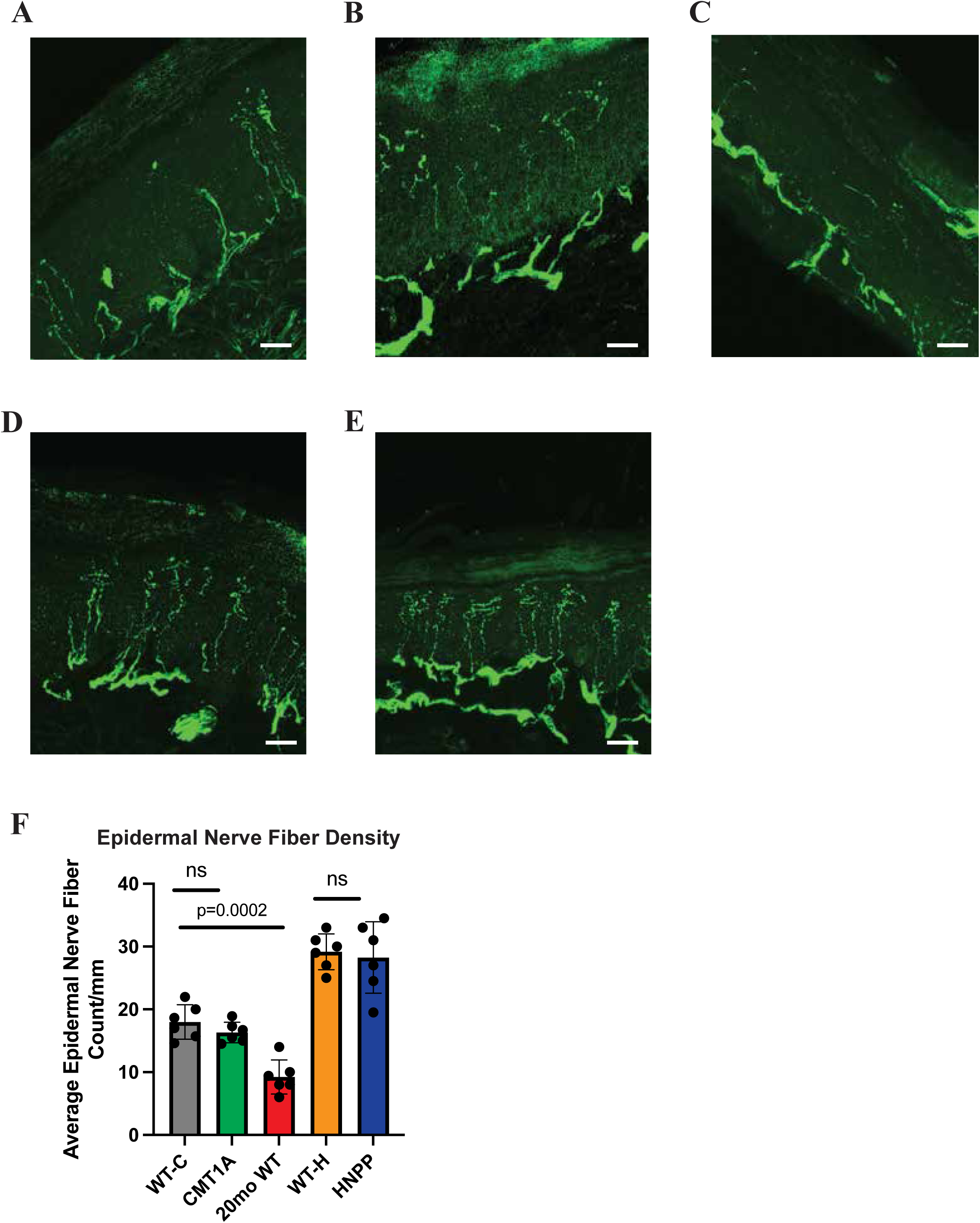
Reproducibility of thermal gradient and hot plate testing. (A) Temperature and time distribution across runs of the thermal gradient assay. (B) Overall temperature preference between runs. (C) Distance and time distribution along the gradient, and (D) total distance traveled between runs. (E–F) Intra- and inter-rater reliability for hot plate testing. n = 10 total animals (5 WT-C, 5 CMT1A). One-way ANOVA followed by Bonferroni post hoc test was used for panels A and B; Student’s t-test for panels C and D. Pearson correlation coefficients (p-values shown) were calculated for panels E and F. Data in panels A-D are represented as the mean ± SEM, mean data points for each animal are shown in panels E,F.

Hot plate testing demonstrated moderately strong intra-rater reliability, supported by Pearson correlation coefficients (Figure 4E). Inter-rater reliability was lower but still demonstrated a borderline significance (Figure 4F). Importantly, genotype-dependent differences between wild-type and CMT1A mice were consistently reproduced, indicating robust biological reproducibility despite operator variability.

### 3.5 Intraepidermal Nerve Fiber Densities (IENFD)

We next quantified intraepidermal nerve fiber density in paw pads. CMT1A mice exhibited preserved epidermal innervation, consistent with human CMT1A findings (Figure 5A–B,F). As expected, aged mice showed significant reductions in epidermal fibers, consistent with age-related small-fiber degeneration reported in humans. Both WT-H and HNPP mice exhibited similar IENFDs (Figure 5D–F), suggesting that their lack of a thermoffensive behavior was not due to overt small-fiber loss. There was no correlation with IEFND and hot plate testing or temperature preferences.

**Figure 5.**
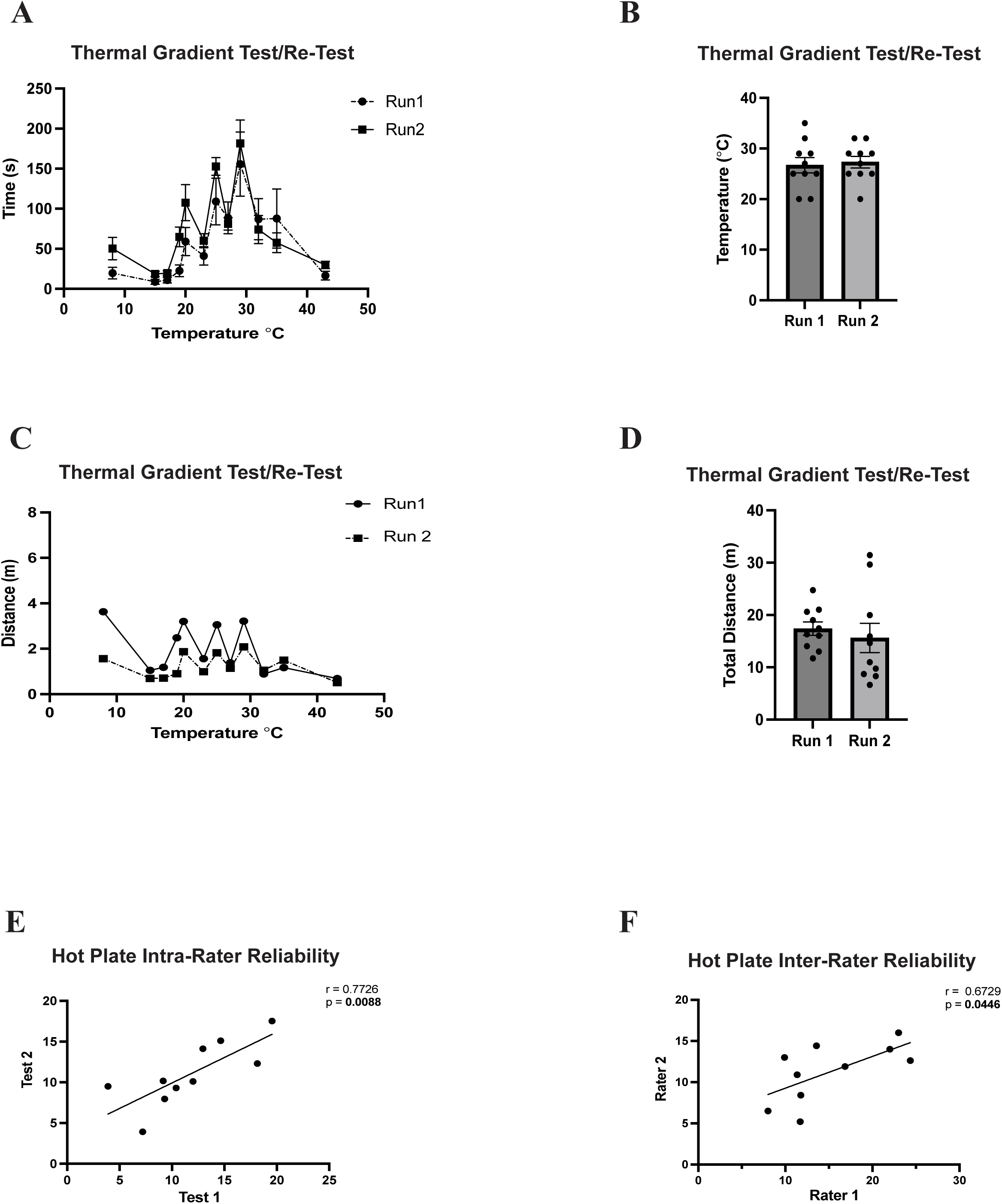
Intraepidermal nerve fiber density across genotypes. Representative images of paw pad skin biopsies stained for PGP9.5 (green) in (A) wild-type, (B) CMT1A, (C) 20-month-old wild-type, and (D) HNPP mice. (E) Quantification of total fibers per mm in each animal. n = 6 per phenotype; data represent the average of three 1-mm segments per animal, group mean ± SEM is represented on graph. Statistical differences were analyzed using Student’s t-test.

### 3.6 TRPV1-Positive Neurons in DRGs: Quantification and Morphology

Given the limitations of TRPV1 immunostaining in skin and nerve cross-sections—where strong epithelial expression can obscure TRPV1 signals within axons, we focused our analyses on dorsal root ganglia (DRG) to better resolve morphological changes in TRPV1-positive neurons that may underlie the thermoceptive dysfunction observed in aged and CMT1A mice. Because thermoceptive deficits were not clearly identified in HNPP mice, subsequent analyses were restricted to aged and CMT1A groups. Co-labeling with TRKA and NEFH was used to distinguish TRPV1-positive Aδ-fibers (TRKA⁺/NEFH⁺) from C-fibers (NEFH⁻ or NEFH^low^). Representative images for each phenotype are shown in Figure 6A.

**Figure 6.**
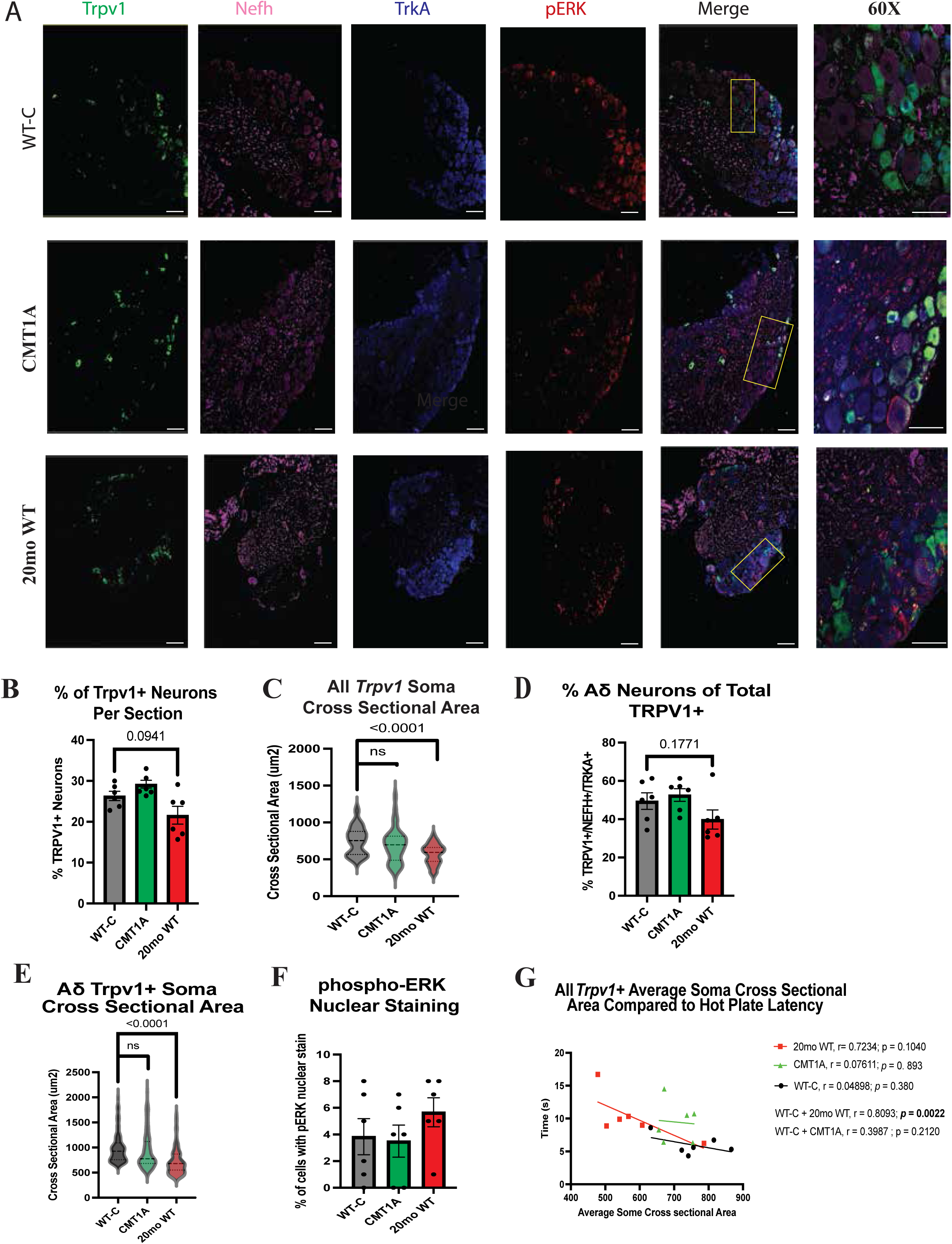
Morphological and molecular characterization of TRPV1⁺ DRG neurons. (A) Representative DRG immunostaining from wild-type, CMT1A, and 20-month-old mice. (B) Percentage of TRPV1⁺ neurons relative to total neurons counted in DIC widefield images. (C) Soma cross-sectional area of the total TRPV1⁺ population (n = 6 animals per phenotype; neurons measured: WT-C = 187, CMT1A = 191, 20mo WT = 185). (D) Percentage of TRPV1⁺/NEFH⁺/TRKA⁺ neurons per phenotype (neurons counted: WT-C = 77, CMT1A = 75, 20mo WT = 75). (E) Soma cross-sectional area of TRPV1⁺/NEFH⁺/TRKA⁺ neurons. (F) Percentage of all neurons staining for phospho-ERK1/2 (n = 100 neurons per animal per phenotype). (G) Correlation between hot plate latency and soma cross-sectional area of TRPV1⁺ neurons (n = 6 per phenotype). Statistical comparisons among phenotypes (B–E) were performed using Student’s t-test; Pearson correlation coefficients (G) were calculated for each phenotype and combined datasets.

Both aged and CMT1A mice exhibited a trend toward an increased proportion of TRPV1-positive neurons compared with young wild-type controls, although these differences did not reach statistical significance (Figure 6B). Soma-size analysis revealed significantly smaller TRPV1-positive neurons in aged mice relative to young controls, whereas CMT1A mice showed no such difference (Figure 6C). Further, aged mice demonstrated a shift toward smaller-diameter Aδ-fiber populations (TRPV1⁺/NEFH⁺/TRKA⁺), consistent with selective vulnerability of Aδ-neurons (Figure 6D–E), while no comparable shift was observed in CMT1A mice. Phospho-ERK1/2 immunostaining indicated uniformly low basal activation across all groups, suggesting no genotype- or age-dependent differences in baseline nociceptive signaling (Figure 6F).

In a subset of animals, we compared hot-plate latency with the mean soma cross-sectional area of total TRPV1⁺ and TRPV1⁺/NEFH⁺/TRKA⁺ populations. Across groups, smaller TRPV1⁺ soma size tended to associate with longer hot-plate latencies (Figure 6G). When data were combined for WT-C + CMT1A and WT-C + aged wild-type groups, a strong correlation between TRPV1⁺ soma size and hot-plate latency was observed in the aged cohort but not in CMT1A mice. These findings suggest that thermal deficits in older animals likely stem from neuronal or axonal dysfunction, whereas prolonged response latencies in CMT1A mice arise primarily from dysmyelination. No significant correlations were detected between TRPV1⁺/NEFH⁺/TRKA⁺ populations and hot-plate or thermal preference outcomes.

## DISCUSSION

In this study, we systematically characterized thermoceptive behaviors in mouse models of two common inherited dysmyelinating neuropathies (C3-PMP22 and LacZ-HNPP) as well as in naturally aged mice. Despite their distinct underlying pathophysiologic mechanisms, aberrant thermoception is a prominent clinical feature in both inherited neuropathies and aging. Yet, experimental investigations of small-fiber function have been limited, as most prior studies have focused on large soma diameter modalities such as mechanosensations. Our findings demonstrate that both aging and dysmyelinating neuropathies impair thermoception through divergent mechanisms.

Characterizing sensory responses in animal models presents inherent challenges, as assays generally assess maximal thresholds (e.g., pressure, heat, cold) rather than the sustained, low threshold activity of C-fibers. Thermal gradient assays offer a unique advantage because they provide a non-forced, user-independent measure of thermal preference across the physiological range. This approach, validated in diabetic neuropathy models (*22*), allowed us to examine subtle behavioral changes in the present study. We found that aged animals, but not CMT1A or HNPP mice, displayed a preference for cooler zones, consistent with TRPV1 ⁺ C-fiber impairment and reduced intraepidermal fiber density. These findings parallel human data showing age-related loss of small fibers and reduced responses to innocuous thermal stimuli. However, thermal gradient behaviors may be influenced by multiple factors such as locomotor endurance and exploratory drive, as evidenced by the reduced total travel distances in aged and HNPP mice. Thus, while the thermal gradient assay showed strong reproducibility, activity-related metrics should be interpreted cautiously and supplemented with complementary behavioral tests.

When examining Aδ fiber-mediated responses using hot plate testing, we observed significant differences in both CMT1A and aged mice. Although delayed hot plate response in CMT1A mice have been reported previously, the prolonged interval to response of greater than 60 seconds is likely beyond Aδ conduction alone (*23*) or representing a plate temperatures below 51-53°C. In this study the elevated thermal thresholds in both CMT1A and aged mice, reflected by prolonged nocifensive responses, were consistent with impaired Aδ-mediated nociception given the rapid responses in young wild type mice. This finding aligns with clinical finding in CMT1A patients, which Aδ-fiber dysfunction is strongly associated with neuropathic pain despite preserved intraepidermal nerve densities. Interestingly, HNPP mice on the 129S background did not exhibit consistent nocifensive responses, even at noxious temperatures, suggesting the possibility that thermoceptive dysfunction in this model may require either strain-specific factors or mechanical injury to manifest. These findings emphasize that genetic background and experimental context critically shape behavioral phenotypes.

The reproducibility of behavioral assays is crucial for translational relevance. We found that the thermal gradient paradigm showed high intra-rater reproducibility across repeated trials, consistent with a user-independent system design. In contrast, hot plate testing showed modest intra-rater but weaker inter-rater reliability, underscoring the importance of consistent operator performance in longitudinal studies. Despite this limitation, genotype-specific differences were reliably detected, suggesting that hot plate testing remains valid approach when it is carefully standardized.

To determine the morphological and structural correlates of behavioral deficits, we examined intraepidermal nerve fiber densities (IENFD) and DRG based on previous finding of an age-related shift towards a higher number of small diameter neurons (*24, 25*). As expected, aged mice exhibited a significant epidermal fiber loss, which is consistent with altered thermal preference behaviors. In contrast, CMT1A and HNPP mice displayed preserved epidermal fiber counts, consistent with what is reported in humans, suggesting functional rather than structural small fiber abnormalities. Analysis of DRG immunostaining further clarified these differences. In aged mice, TRPV1⁺ neurons were smaller soma size, particularly within the TRPV1⁺/NEFH⁺/TRKA⁺ Aδ population, consistent with a degenerative or senescent shift, which may underlie impaired Aδ-fiber responses. In contrast, TRPV1⁺ neurons in CMT1A mice displayed preserved morphology, indicating that functional deficits likely stem from dysmyelination and slow conduction rather than axonal degeneration. These findings highlight distinct mechanistic pathways by which aging and inherited neuropathies disrupt thermoception.

An additional consideration is the paradoxical relationship between reduced thermal perception and neuropathic pain. Many patients with inherited neuropathies or advanced age report burning dysesthesias(*26*), suggesting cycles of hypo- and hyper-activation within dysfunctional sensory neurons(*27*). To address whether basal neuronal hyperactivation contributes to this phenomenon, we evaluated phospho-ERK1/2, a marker of nociceptive pathways, and detected uniformly low expression levels across all groups. This finding were consistent with the absence of baseline hyperactivity. Future studies should assess additional signaling markers such as p38 MAPK and evaluate responses following thermal challenge to determine whether pathological signaling emerges under stress conditions.

Correlation analyses provided further evidence linking morphological and functional changes. Across animals, smaller TRPV1⁺ neuron soma area correlated with longer hot-plate latency times, suggesting that neuronal atrophy contributes directly to delayed nocifensive responses. This relationship was significant in aged mice but not in CMT1A, reinforcing the concept that neuronal degeneration, rather than dysmyelination, underlies thermal deficits in aging, whereas impaired conduction dominates in CMT1A.

## CONCLUSION

In this study, we characterized thermoceptive behaviors and associated structural changes in epidermal axons and DRG neurons across models of inherited dysmyelinating neuropathy and natural aging. CMT1A and aged mice recapitulated key thermoceptive deficits observed in patients, whereas HNPP mice exhibited less consistent phenotypes. Aging was associated with reduced IENFD and morphological alterations in TRPV1⁺ Aδ neurons, consistent with degenerative or senescent processes. In contrast, CMT1A mice displayed preserved neuronal morphology, indicating that thermoceptive impairment arises primarily from dysmyelination and slowed conduction.

Cross-correlation between TRPV1⁺ neuronal soma area and hot-plate latency revealed a strong inverse relationship in aged, but not CMT1A, mice. This observation underscores the mechanistic divergence between neuronal morphologic associated thermal loss in aging and conduction related delay in dysmyelination, offering a potential quantitative biomarker for distinguishing these processes in future translational studies.

Both thermal gradient and hot plate assays demonstrated strong reproducibility and biological sensitivity, supporting their utility as preclinical outcome measures for small-fiber dysfunction. Together, these findings highlight thermoceptive impairment as a shared manifestation of aging and inherited neuropathies arising from distinct mechanisms. Therapeutic strategies should therefore be tailored to these underlying differences. By validating behavioral and morphological endpoints, this study establishes a framework for translational investigations targeting thermoceptive dysfunction and neuropathic pain in peripheral nerve disease.

